# Annotating opportunistic camera-trap images with conditions of recording, for the disease surveillance of Eurasian lynx (*Lynx lynx*)

**DOI:** 10.1101/2025.05.15.654015

**Authors:** Louison Blin, Anouk Decors, Delphine Chenesseau, Laura Lenglin, Marie-Pierre Ryser-Degiorgis, Fridolin Zimmermann, Stéphanie Borel, Suzanne Bastian

**Affiliations:** Office français de la biodiversité (OFB), Direction Régionale-Bourgogne-Franche-Comté, Réseau Loup-Lynx, Dijon, France; Office français de la biodiversité (OFB), Direction de le Recherche et de l’Appui Scientifique, SantéAgri, Orléans, France; Oniris, INRAE, BIOEPAR, 44300 Nantes, France; Institute for Fish and Wildlife Health, Vetsuisse Faculty, University of Bern, Bern, Switzerland; Foundation KORA, Ittigen, Switzerland; University of Lausanne, Department of Ecology and Evolution, Lausanne, Switzerland

**Keywords:** Camera trap, Syndromic surveillance, Eurasian lynx, Image quality, Interdisciplinarity

## Abstract

The French population of the Eurasian lynx (*Lynx lynx*) is small and fragmented. Any emerging disease would endanger it even further, so health surveillance is crucial. Currently, health monitoring relies on lynx carcass surveillance. In parallel, the Eurasian lynx population is being monitored since 1997 through a large network of observers in different regions and this trove of camera-trap images could allow for the opportunistic detection of clinical signs. Camera traps have been used for a very long time in ecology and, more recently, in epidemiology to study *e*.*g*. sarcoptic mange. However, the quality of the images from camera traps varies, the details of the animal’s body are more or less clearly visible. This work examines how the quality of the images relates to the ability to detect cutaneous changes and abnormal body conditions. Different factors affect image quality and the detection of changes: intrinsic camera parameters like the type and settings of the camera trap, extrinsic factors like the external lighting conditions or the position of the animal in relation to the camera. In our data set, clearly visible cutaneous changes were associated with a different set of factors than visible abnormal body conditions. The camera-trap conditions currently used for ecological monitoring of the lynx are sufficiently diverse to allow for the general surveillance of abnormal health signs. However, for monitoring specific health signs, the camera settings as well as the shooting context should be optimized to ensure the best possible sensitivity and specificity of the detection.

## 1 Introduction

The Eurasian lynx (*Lynx lynx*) is a species with a very wide range, from the Palaearctic regions to western Europe and eastern Asia (Pascal et al., 2003). There are eleven isolated populations of lynx in Europe (Kaczensky et al., 2021), three of which are found in France: the Franco-Swiss population in the Jura Mountains, the Franco-German population in the Vosges-Palatinian and the Alpine population (Kaczensky et al., 2013). These populations are all the result of reintroduction programmes initiated in the 1970s (Chapron et al., 2014), with individuals mainly coming from the Carpathian population (Mueller et al., 2022). In France, populations are stable or increasing but remain threatened (Kaczensky et al., 2013; Office Français de la Biodiversité, 2022). The species is classified as of Least Concern by the IUCN at a global level, but locally, the Alps and Jura populations are considered as Endangered and the Vosges-Palatinian population as Critically Endangered (von Arx 2020). The main threats to these populations are illegal killing, inbreeding, habitat fragmentation and mortality due to collisions with vehicles (Kaczensky et al., 2013). In France, the latter cause appears to be the main threat to the species (Drouet-Hoguet et al., 2021), but roadkill is also the most frequent way of detecting lynx carcasses and mortality due to diseases could be underestimated (Schmidt-Posthaus et al., 2002). It remains that in such fragmented and fragile populations, the epidemic spread of a disease can represent a very serious threat. Even with a low mortality rate, in synergy with other mechanisms, a change in the population’s behaviour or its demographic parameters can precipitate the extinction of endangered populations (Preece et al., 2017). Lynx are difficult to monitor because they are generally nocturnal and secretive. They have large home ranges, often live in dense cover habitat and occur at low population densities. In France, until 2020, their health surveillance was mainly based only on necropsies. However, this method is hampered by the scarcity of carcasses typical for lynx which occur at low densities and arenot hunted (Heiderich et al., 2023). Camera traps are being used more and more often to monitor diseases in wildlife, particularly in carnivores. Studies have used them for monitoring sarcoptic mange in red foxes (*Vulpes vulpes*) (Barroso and Palencia, 2024; Carricondo-Sanchez et al., 2017; Pisano et al., 2019), in grey wolves (*Canis lupus*) (Oleaga et al., 2011) and in coyotes (*Canis latrans*) (Murray et al., 2021). Muneza et al. (2019) used them to monitor and characterize limb lesions on giraffes (*Giraffa camelopardalis*).

The population monitoring of the Eurasian lynx in France is done by the *Réseau Loup-Lynx* - Wolf-Lynx Network - run by the French Agency for Biodiversity (OFB). The aim of the network is to obtain robust and reliable information on their spatial distribution, their numbers and their damages to livestock, to better inform political decisions and management measures. The data comes from chance observations and opportunistic camera trapping. The identification of individuals provides more information on the biology, ecology and monitoring of lynx (minimum number of individuals, information on roaming area, livestock raiders, pattern of sub-adults dispersal, number of breeding females incl. juveniles, period of presence of an individual or to identify its mother, which are important pieces in the kinship analyses, etc.) (Gimenez et al., 2019; KORA Foundation, 2022). In France, all the images are centralised and annotated by two specialists from the OFB and stored in the national photo-identification database.

There are also specific camera trapping campaigns in some areas, but not every year. Some campaigns on the Franco-Swiss population in the Jura Mountains were performed jointly by the OFB in France and KORA in Switzerland. KORA, the foundation for Carnivore Ecology and Wildlife Management, monitors large carnivores on behalf of the Swiss Federal Office for the Environment, in close cooperation with the cantons. In the specific campaigns, the photo-identification is also used to estimate lynx densities by means of CR models (Fridolin Zimmermann et al. 2013; F Zimmermann, Foresti, and Zimmermann 2016).

We sought to evaluate how the population monitoring images resulting from camera traps could complement an event-based health surveillance that is based on pathological examinations of carcasses. For this, two questions were addressed in parallel studies: here, we examine the features of images that allow the detection of clinical signs. Another paper documents the variety of clinical signs that have been observed in the national database and their relation to disease and injuries of the lynx (Lenglin et al., submitted 2025).

In this paper, we sought to contrast the features of images where signs are clearly visible with images where clinical signs were not visible or of low confidence. Ideally, we should have compared pairs of images from the same events, with or without clearly visible clinical signs, but there were not enough images to do that. We chose an exploratory strategy of a statistical association between features and clinical sign detection, on a convenience sample. We did not aim at comparing images for diseased animals versus healthy animals (this was not a case-control study).

## 2 Materials and Methods

### 1.1 Data collection

The OFB has been organizing the monitoring of the Eurasian Lynx population since 1997, across its area of distribution, in mainland France. A network of field observers, most professional, some amateur, provide images and videos from camera traps to regional network coordinators. The latter sorts the images, identify the individuals by their fur markings if image quality allows it, then store the files in a database grouped as « events ». An event is made of one or several detections of the same animal, over a period of 12 hours. Sometimes cubs (non-identified) are recorded with their mother in the same event. If two animals are present and identifiable, 2 events are created.

For identification, the recommended setting of camera traps is along a path that the animal uses regularly. When possible, two cameras are placed on either side of the path so they can capture both flanks. In this study, this setup is called “on-trail”. Sometimes, when a lynx kill is detected in the field, a camera trap is set up near the carcass to photograph the animal when it comes back to feed. We call this setup on “prey”. The picture is then taken from various angles, depending on the movements of the animal.

### 1.2 Sampling design

We selected a subset of the images of the database with the following steps, as illustrated in Figure 1:

**Figure 1.**
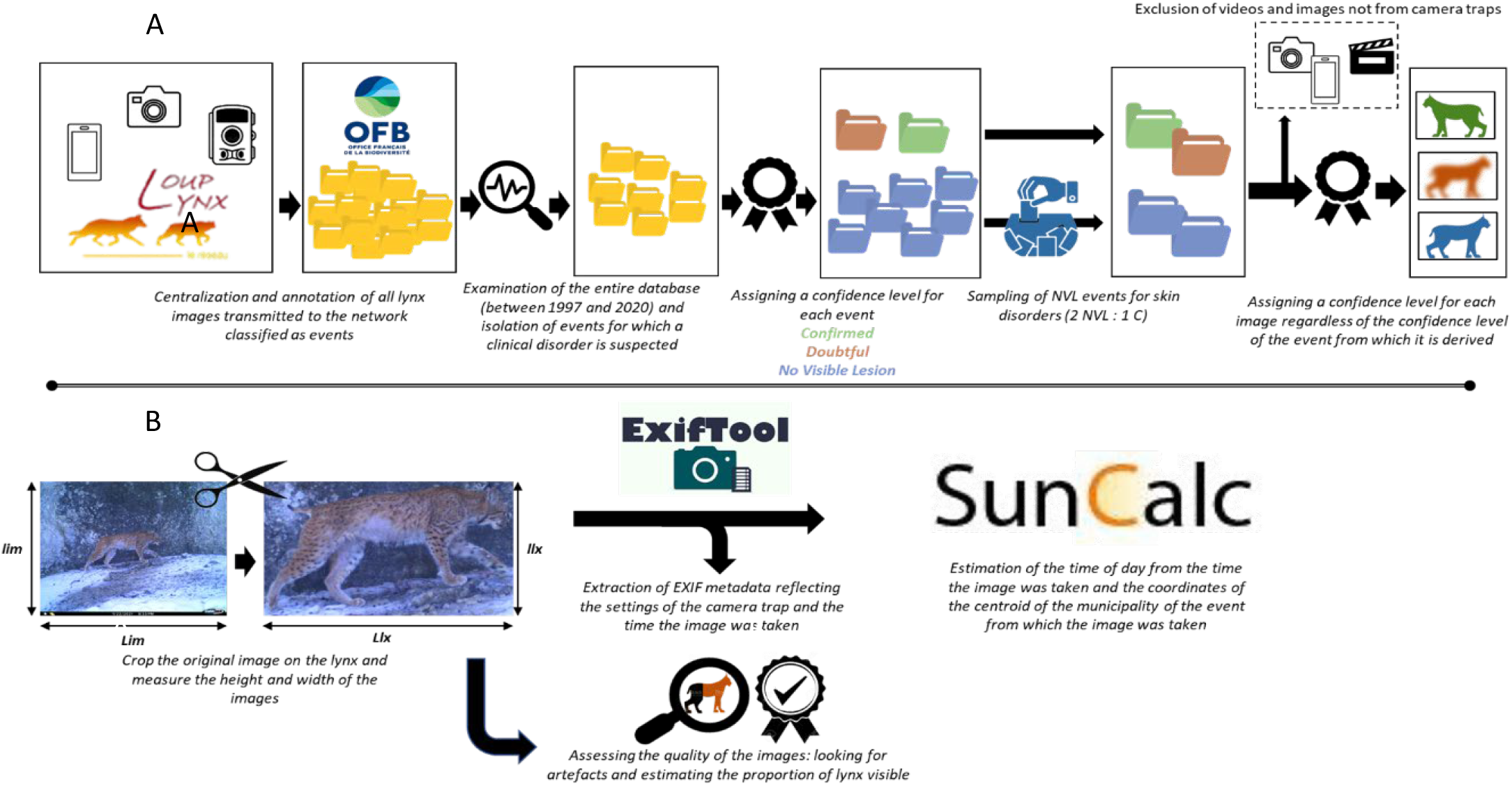
Summary of steps carried out from data acquisition (A) to image quality annotation and features calculation (B).

- firstly, two veterinary students without an experience of disease detection browsed the whole database (4035 events, 29974 images). Videos and images from other kinds of cameras (phone, camera) were not included in the present study. In total, they flagged 638 events captured by camera-traps as suspect of showing clinical abnormalities in at least one image. The median of the number of images per event was 2 for « on-trail » events and 20 for « on prey » events.
- secondly, these flagged events were reviewed by a veterinarian wildlife expert, who categorized 90 of them as « confirmed » if a sign was clearly visible on one or more images or “doubtful”, if only one image had possible signs that were unclear. After this, 548 events out of 638 had in fact “no visible pathological changes” (code “NPC”) and 27 were “doubtful”. Among the 63 “confirmed” events, two kinds of clinical signs were most represented: skin lesions and abnormal body conditions. Some animals showed several clinical signs simultaneously, *e. g*. alopecia and cachexia.

The next steps aimed at selecting a set of images, such as to compare the features of images where clinical signs are clearly visible with those where they are not. We selected only categories for which a high number of confirmed events were available: skin (cutaneous) lesions called “cutaneous changes” and abnormal body conditions called “body condition changes”.

- the expert veterinarian qualified each image inside the “confirmed” or “doubtful” events, as “clearly visible CV”, “low confidence LC” or “no visible pathological changes NPC”.
- we built the convenience sample of contrasted images, by selecting all images from the 63 “confirmed” events (all their CV, LC and NPC images) and from the 27 “doubtful “events (LC and NPC images). To have enough NPC images, we completed this set with a subset of images from “NPC” events, such as to balance the number of images taken from prey and from on-trail events and to have a ratio of between 2 and 4 « NPC » images for each « clearly visible » image. We discarded images where we did not have metadata of image features.

### 1.3 Co-variates

We annotated the images of this reduced dataset with 4 sets of variables:

#### 1.3.1 Camera settings

The model of the camera; the exposure value as a measure of the quantity of light entering the camera, in given conditions of lighting conditions; the diaphragm aperture; the diaphragm shutter speed as a measure of exposure time; the light sensitivity or “ISO” value; with or without flashlight; the color range of the image *i*.*e*., “black and white”, “pink and white” or “color”. These information were extracted from the EXIF metadata of the image files, with help of the ExifToolR package (O’Brien, 2024) under R (R Core Team, 2022).

#### 1.3.2 The external conditions of visibility

We calculated intervals of lighting conditions (the position of the sun in the sky) as a function of date, time and geographical location of the image, with help of the package *Suncalc* (Thieurmel and Elmarhraoui, 2022). We classified images between dawn and dusk as « Daytime » and between dusk and down as « Nighttime ». For most images, the precise GPS location was not available in the metadata, so we approximated it by calculating the centroid of the district address where they were taken (the French administrative “commune” has an average area of 14 km^2^) with the *sf* package (Pebesma et al., 2023). The weather may influence also the external conditions of visibility, but the meteorological data that were available from regional stations were only provided in 8-hour intervals, at a distance of up to 100 km, and at a lower altitude than most lynx observations, so we did not use them.

#### 1.3.3 The distance, angle and proportion of visible parts of the animal on the image

We annotated each image manually with a list of visible parts of the body (back, front, left flank etc.) and used this information to calculate the proportion of the body that is visible, between 0.25 (¼^th^ of the animal) and 1 (whole animal) (Supplementary materials 1). We calculated an indicator of the distance to the camera, by comparing the dimensions in pixels of the image cropped to the visible body parts of the animal with the dimension of the entire image. The ratio was weighted with the proportion of visible body parts, to simulate how big the cropped image would be if the whole animal was visible.

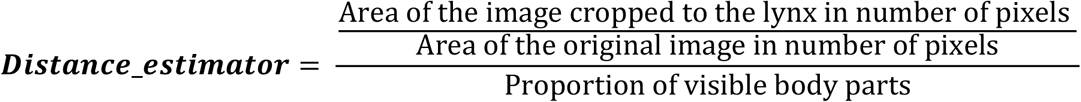

The higher this indicator, the closer the lynx was to the camera trap. The reference value of 1 means that the whole flank of the lynx is visible and that it covers the entire image.

As an indicator of image ***definition***, we used the area *i*.*e*. the number of pixels of the rectangle surrounding the cropped image of the lynx.

From the list of visible body parts, we also created the variable ***Profile***, with four categories:

- **Back**: if the lynx was perpendicular to the camera trap and the tail and the back of the ears were visible;
- **Face**: if the lynx was perpendicular to the camera trap and the head was visible (right hemiface (called after right face) and left hemiface (called after left face);
- **Right_Profile**: if the lynx was parallel to the camera trap and the right profile was visible
- **Left_Profile**: if the lynx was parallel to the camera trap and the left profile was visible

Then, we measured the quantity of visible profile translated by the variable

***Profile_completeness*** (Supplementary Material 2) as follows:

- For profiles **Face** and **Back, *Profile_completeness*** took the value **0.25** by default.
- For **Right_** and **Left_Profile**: we looked at how much of the profile was visible: o the entire profile was visible, ***Profile_completeness*** took the value **1**
  ∘ three quarters of the profile were visible, ***Profile_completeness*** is set to **0.75**
  ∘ half of the profile was visible, ***Profile_completeness*** is set to **0.5**
  ∘ a quarter of the profile was visible, ***Profile_completeness*** is set to **0.25**

#### 1.3.4 Visible artifacts

We manually annotated each image with identifiable artifacts of the following categories:

- The presence of a blur in the image (***blur***)
- The presence of over- or underexposed areas in the image (***presence of light artifacts***)
- The presence of vegetation obstructing all or part of the lynx (***presence of environmental artifacts***)
- The presence of elements related to weather conditions, *e*.*g*. snowflakes, likely to interfere with a clear understanding of the image (***presence of weather artifacts***)

### 1.4 Statistical analysis

In this dataset, we calculated the pair-wise association between each variable and the detection of clinical signs: cutaneous changes on the one hand and body condition changes on the other, each with two modalities (Yes/No). When the explanatory variable was quantitative, we used a Wilcoxon-Mann-Whitney test to compare distributions. When the explanatory variable was qualitative, we used either a Chi-square test or a Fisher’s exact test.

Secondly, we explored the association between groups of variables and the detection of skin changes on the one hand and body state changes on the other. We used multivariate analyses (Principal Component Analyses (PCA) and Mixed Component Analyses (MCA)). The analyses were performed with the *FactoMineR* package (Husson et al., 2023) interfaced with the *FactoShiny* package (Vaissie, Monge, and Husson 2021). Graphical representations were produced using the ggplot2 package (Wickham, 2016).

All operations were performed using R software (R Core Team, 2022) interfaced with RStudio (RStudio Team, 2020).

## 2 Results

The list of visible clinical signs and their diagnostic significance are detailed in a parallel study (Lenglin et al., submitted 2025). Here we look at the general quality of the images and the conditions of shooting, in a subset of 144 images, of which 22 had “clearly visible” cutaneous signs, 12 had “clearly visible” abnormal body condition signs.

No information about the type of flash is available in the EXIF metadata, so the only information that allows us to approximate the type of flash is the type of image produced: colour or not. In our dataset, 86.1% of the images were colour images (19 black and white photos and 1 photo in shades of pink), i.e. from a white flash (Xenon or white LED). In our dataset, 23 images were from events obtained by camera traps set near prey (prey image), so 84.0% of the images were on-trail images.

### 2.1 Diversity of cameras and settings in the dataset

The 144 images that were used in this study came from 18 models of camera traps belonging to 7 brands, most of them (115 images) of the *Cuddeback* brand, the most frequent being *Attack* (34 images), *Ambush* (30 images) and *C1* (20 images).

The relative settings of these camera traps are automatic and depend on external lighting conditions. Intrinsic parameters like the exposure value, the aperture, the shutter speed or the ISO sensitivity are mutually dependent and cannot be modified by the operator in these devices. The only parameter that can be controlled by the operator is the triggering of the flash and possibly its intensity. As expected, a PCA comparing the settings between images showed a collinearity between the exposure value and the shutter speed, both independent from the aperture and the ISO level, which were inversely colinear. In order to reduce the number of variables, the settings parameters were thus reduced to the exposure value and the ISO level.

### 2.2 Correlation between the detection of clinical signs, external conditions of lighting and camera settings

#### 2.2.1 External conditions of lighting, camera settings and detection of cutaneous changes

The images were taken at different times of day and during different months throughout the years (Figure 2).

**Figure 2.**
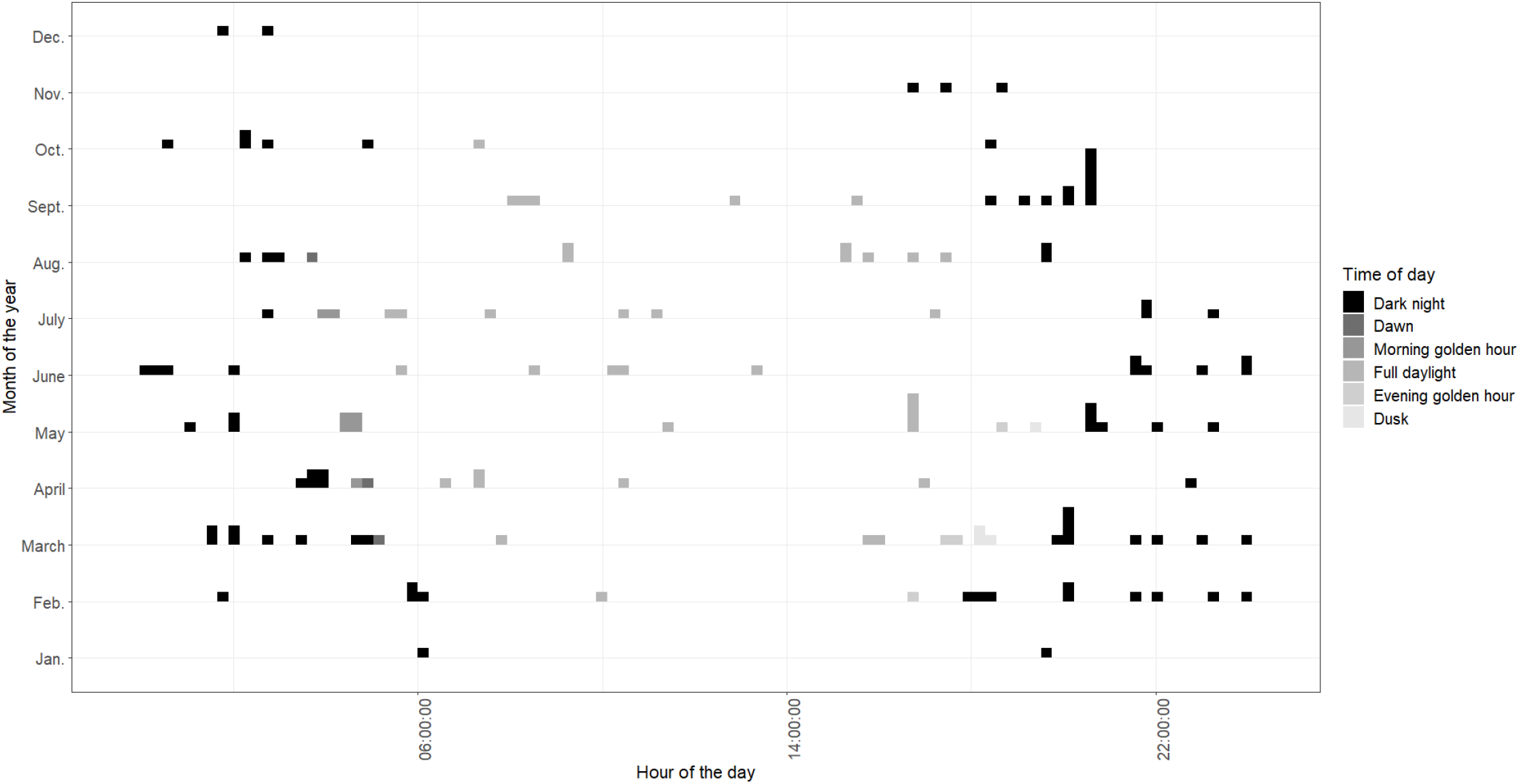
Diversity of times and days of capture in the dataset. The figure shows the number of images taken per hour for each month of the year. The corresponding time of day is indicated by the colors. The times of day corresponding to a nocturnal lighting atmosphere (**DarkNight, Dawn** and **Morning golden hour** modes). The times of day corresponding to a diurnal lighting atmosphere (Full **Daylight, Evening golden hour** and **Dusk** modes).

With a PCA, the detection of skin changes was associated with taking images at night, *i*.*e*., with low ambient luminosity (low exposure value) and a white flash, to generate color images. The ISO level did not seem to be associated with the detection of cutaneous changes, probably because the flash makes ISO adjustment unnecessary.

#### 2.2.2 External conditions of lighting, camera settings and detection of body condition changes

The detection of body condition changes was associated with daytime image capture, in conditions of high luminosity (high exposure value), images in color and the absence of flash triggering, which is logical during daytime.

In a pairwise comparison (Supplementary materials 3), the flash was negatively associated with the detection of body condition alterations (p-value of 0.03 with a chi-square test).

#### 2.2.3 Position of the lynx and detection of lesions

We plotted the ratio of the ***length of cropped image on lynx/length of entire image against the ratio f(height of cropped image on lynx/height of entire image)***, the images fell into 3 main zones, associated with the context in which the image was taken (Figure 3). This graph suggested that in this dataset, animals captured while feeding on a carcass are more frequently distant and either perpendicular or only partially visible, whereas the on-trail setting capture the flanks of individuals and their size on the image is proportional to the distance.

**Figure 3.**
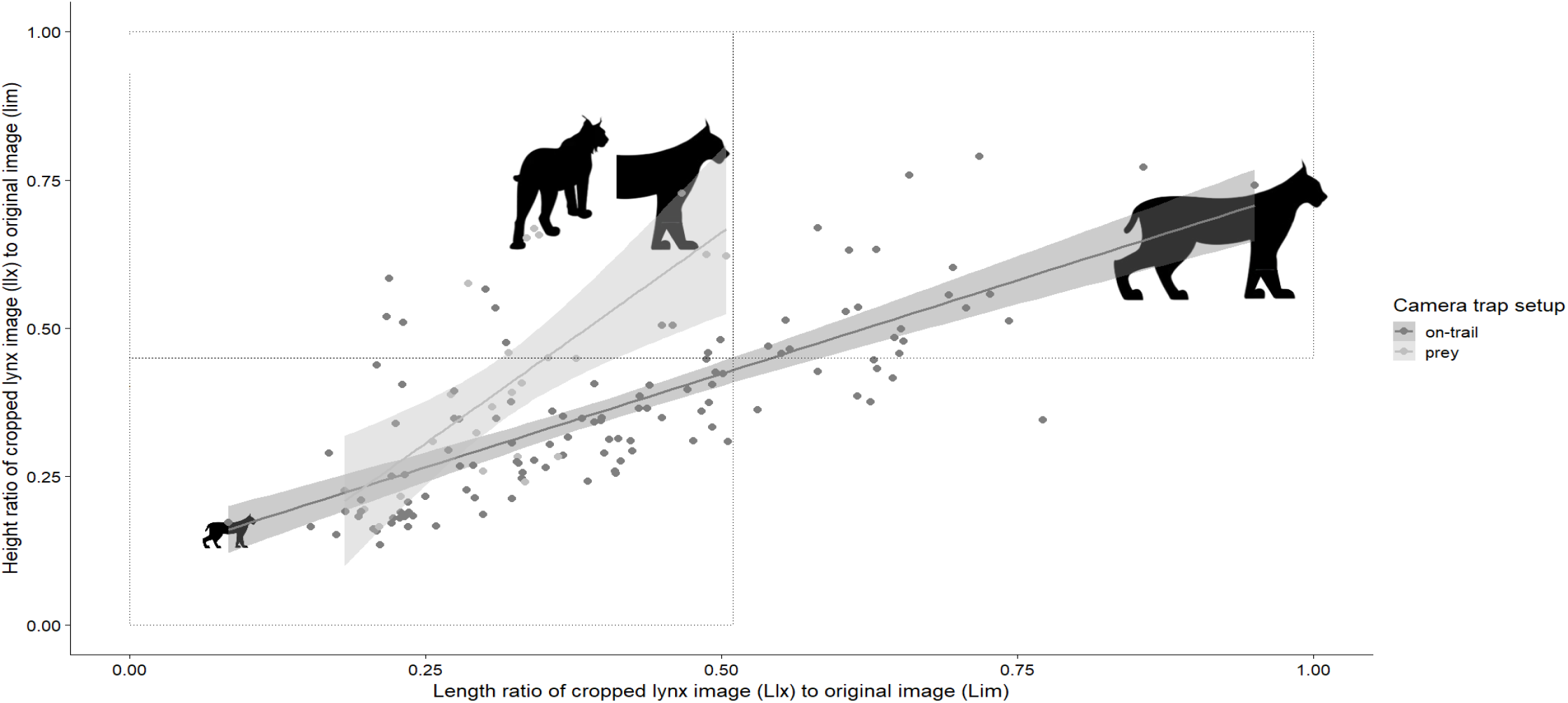
Linear regression of the relative height and length of the cropped images compared to the entire images, in two shooting contexts, on-trail vs. on prey. The silhouettes represent the interpretation regarding the angle and proportion of visible body parts.

##### 2.2.3.1 Lynx position and detection of cutaneous changes

These tendencies are also visible with a PCA analysis (Figure 4). As expected, the “on-trail” setup of cameras was associated with *Right_* and *Left_* flanks that were generally complete (values of 75 to 100%). With prey, the lynx was generally further away (point closer to the lower left corner) from the camera trap and associated with more diverse but less complete profiles, *i*.*e*., between 50 and 75% in a prey context. The *Back* and *Face* profiles visible parts were projected onto the 25% axis, but this is by construction (see Materials and methods).

**Figure 4.**
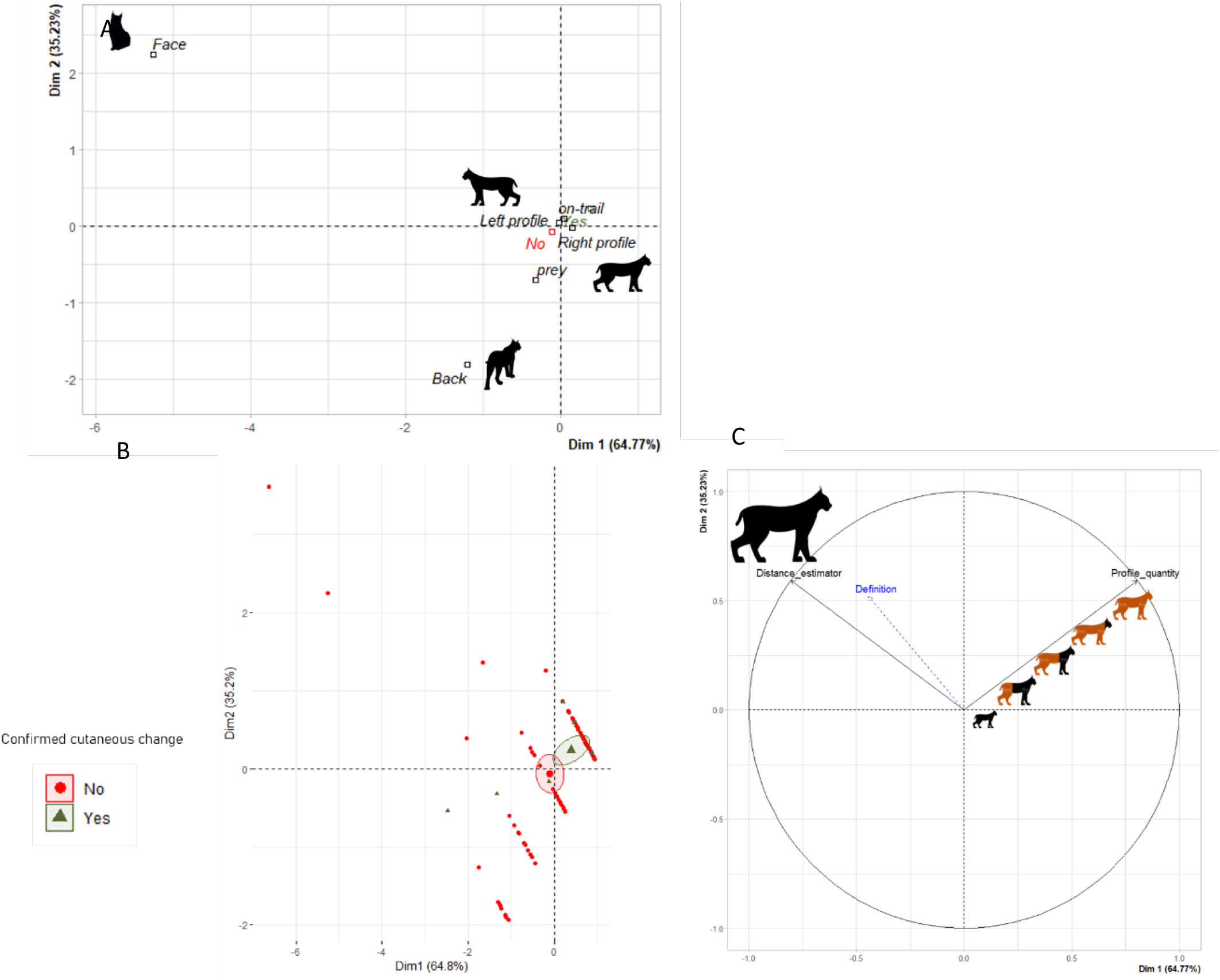
Principal component analysis (PCA) plots illustrating the relationship between the detection of cutaneous changes (“Yes” in green or “No” in red) and two quantitative variables: the estimator of distance to the camera and the profile completeness. Graph A is the graph of individuals with the barycenter of each category. Images with clearly visible cutaneous signs are associated with an on-trail setup, showing lateral profiles (Right_ and Left_profile). Graph C is the graph of quantitative variables, with two projection axes: one for distance and one for profile completeness. Graph B is identical to Graph A, but it shows individual observations and the 95 % confidence ellipses around the barycenter. Graphs B and C reveal an association between profile completeness and the detection of skin changes, independently of distance. With all three graphs, it appears that the detection of skin changes is associated with greater profile completeness, which is in general a right or left profile, more often observed on “on-trail” setup.

The definition of the image varied in the same direction but in an opposite sense as the distance from the camera trap: the farther away the lynx was, the lower the definition.

**Images with clearly visible cutaneous changes were associated with complete profiles, on-trail setup, and at an average distance. Face and Back profiles were not associated with the detection of cutaneous changes**.

##### 2.2.3.2 Detection of body condition alteration

Conversely to cutaneous changes, the images where body condition changes were clearly visible had no particular association with the position of the lynx (angle and distance) nor the camera setup (on-trail or prey).

#### 2.2.4 Artifacts and the detection of lesions

##### 2.2.4.1 Artifacts and detection of cutaneous changes

The detection of cutaneous changes was associated with the absence of blur in the image at 95% (p-value of 0.02 with a chi-square test), itself associated with a nighttime flash. The presence of light artifacts was associated with the distance to the camera trap: the closer the camera trap, the greater the risk of having a light artifact.

In summary, **cutaneous changes were clearly visible in non-blurred images taken with flash, in conjunction with low exposure levels (*i. e*. at night)**.

##### 2.2.4.2 Artifacts and detection of body condition alteration

The detection of body condition alterations was significantly associated with the absence of flash, logically associated with everything that helps to avoid it. Detection was therefore associated with daylight photography (and therefore a high level of exposure) and a lynx that was fairly close to the camera trap. The presence or absence of blur did not seem to be detrimental to the detection of these changes.

#### 2.2.5 A concrete example: differences in image quality leading to differences in confidence levels within the same event

In Figure 5, the two images shown came from the same event (*i.e*. two images of the same individual taken at the same location in less than 12 hours). On the image on the left (A) a “rat tail” *i.e*. tail alopecia was clearly visible, whereas on the right (B) it was not.

**Figure 5.**
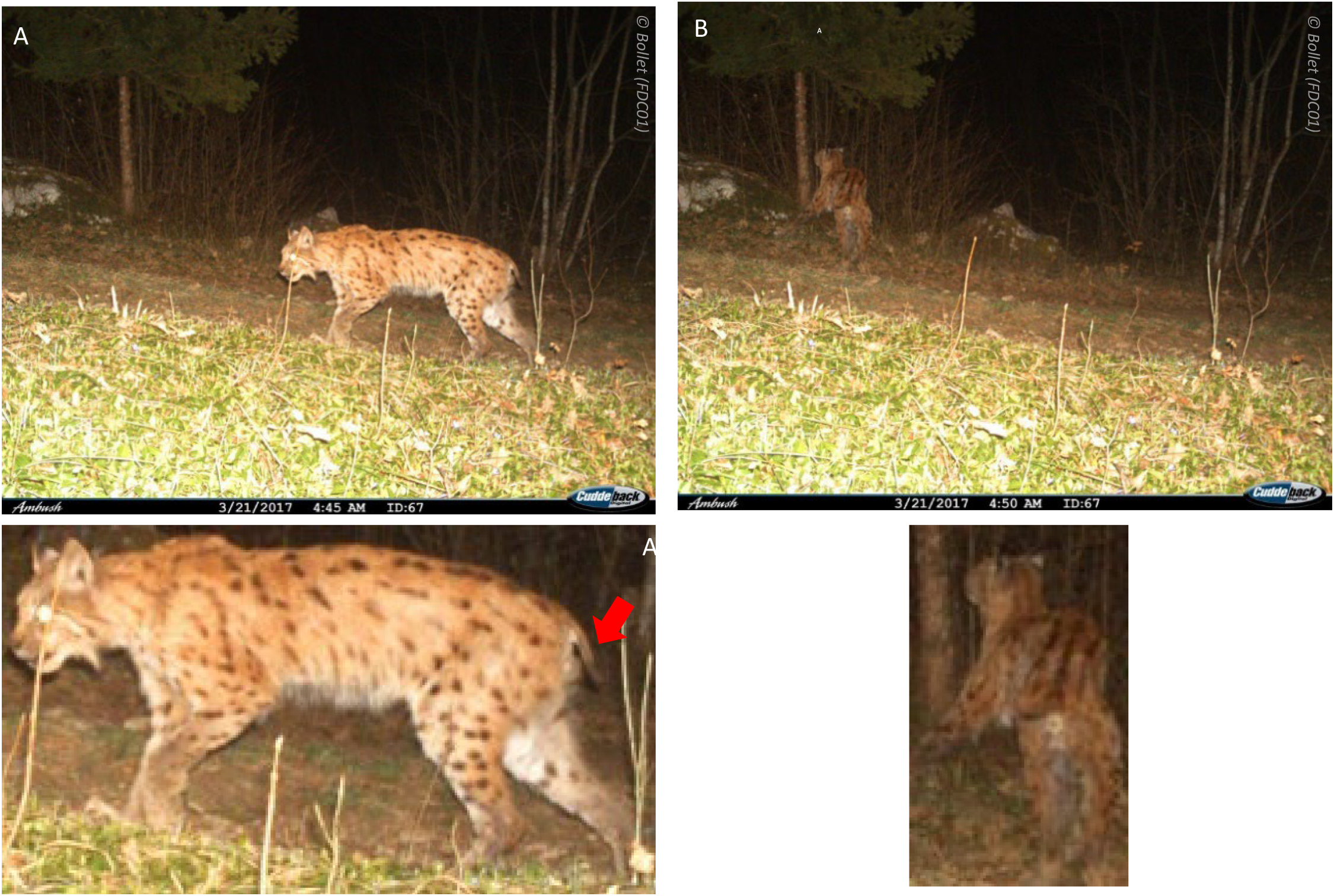
Differences in detection of the “rat tail syndrome” between two images from the same event, with the same camera, in an on-trail setup, at a 5 min interval, on individual F01_042. Images A and A2 show a lynx with clearly visible cutaneous change (identified with the red arrow). Images B and B2 show a lynx with a low confidence cutaneous change. A2 et B2 are images from A and B cropped on the lynx.

In image A, on the other hand, the lynx was closer, the image was better lit and better defined, and the angle of the animal was better suited to detecting the caudal lesion. In image B, the lynx was farther away from the camera trap and there was less light, the image was of poorer definition. In addition, the angle of the lynx was not adequate, the tail was not clearly visible and appears thin, but it was not possible to make an unequivocal decision as to the presence of a lesion.

## 3 Discussion

This protocol aimed at exploring a historical, opportunistic dataset, spanning nearly 30 years of observation, as a resource for a generalist, event-based disease surveillance. The global dataset is made up of images sent to the coordinators of the lynx-wolf network, whose main priority was and remains the photo-identification of individuals for population monitoring.

### 3.1 An exploratory method for examining a set of heterogeneous images

The original database had very heterogeneous photographic methods, settings and capture conditions. It is nearly impossible to have images of reference of diseased lynx taken in different conditions, given their elusive behavior and the rarity of observation. We used a pragmatic approach of screening for sightings with suspected clinical signs, that where further characterized by an expert veterinarian. We then looked for common features in images where signs were clearly visible vs. where they were not. In our subset of images, we found very diverse conditions (setup, type of camera traps, position of the lynx, time of day and year and type of environmental artifacts). We used statistical methods to contrast groups of features, but not to measure the effects of photographic conditions on the sensitivity or specificity of detection, as we have neither a gold standard nor a way to ensure that our subset was representative of the whole database.

This opportunistic data set allowed us to detect a diversity of clinical signs that would not have been visible in a standardized experimental setup. The protocol was very work and time-intensive. In future, our results may be used to train artificial intelligence for screening and annotating images.

### 3.2 Quality of the metadata

In this study we designed an original method for quantifying the position of the lynx in relation to the camera trap. This type of annotation can easily be made manually by delimiting the contours of the lynx (length and height of the lynx) and the visible parts of the lynx to define its distance and angle. Subsequently, this type of operation could also usefully be automated with artificial intelligence. One team has already developed a method and a program on R for measuring animal distances and heights on images from camera traps, without leaving any objects in the camera’s field of view (Strickfaden et al., 2023).

We had no access to sound meteorological or habitat data that would influence lighting conditions and the presence of artifacts. A precise georeferencing of camera-traps and local meteorological measurements could improve our understanding of environmental factors. EXIF metadata is an easily accessible source of information on the settings of the camera and the extraction of this data can be easily automated from the file of the image. However, wildlife cameras optimize the settings automatically, so we in fact observed quite homogeneous conditions minimizing the shutter speed (to avoid the blur) and the aperture (to maximize the depth of field) and those were not distinctive criteria of quality (apart from the flash trigger, which could be set on or off for some models).

In this study, the annotation of the presence of blurs or artifacts (light, environmental, meteorological) was made manually. (Murray et al., 2021) have recently developed a quantitative method to estimate the amount of blur in the image, with grey scale calculations. They defined images as unblurred if their clarity metric was greater than or equal to the mean clarity.

We calculated the position of the sun with help of the *Suncalc* package in R, and we observed that the images from our dataset were taken all day and night. However, there is no benefit in narrowing down this variable to small intervals of time for lighting conditions, as our images clustered into 2 categories: either nighttime or daytime, which results broadly in triggering a flash or not.

### 3.3 Discussion of the results

For detecting cutaneous changes (rat tail syndrome or generalized alopecia), our study suggests that blurs result in a low confidence level or in false positives. For example, a photo of a blurred tail due to movement may appear of a different shape and thickness that falsely suggests an alopecic tail. Authors who studied sarcoptic mange (a disease for which one of the main clinical expressions is the presence of depilated areas, particularly on the tail) in coyotes and foxes have taken into account that blurs are detrimental to detection (Carricondo-Sanchez et al., 2017; Murray et al., 2021). Also, the observation of a fairly complete profile is requested. With on-trail setup, the camera trap is set perpendicular to the path taken by the lynx to view the animal’s flank and/or back, because the animal’s coat patterns are used to photo-identify the individual. This is ideal for detecting skin lesions. Conversely, with prey setup, the position of the lynx is less well controlled because the animal turns around its prey, but at the right distance, it also allows for observations of good quality. A non-perpendicular orientation (face or the back profiles) are suitable to detect clinical changes in other systems, e.g. ocular or genital clinical changes.

We were not able to demonstrate the importance to have a colored image to detect a clinical change in our study but this point is frequently discussed by other authors. Carricondo-Sanchez et al (Carricondo-Sanchez et al., 2017) explained that images taken with infra-red flash, i.e., in black and white, were not suitable for detecting sarcoptic mange. Murray et al. (2021) observed that in blurred black and white images, the probability of detecting a lesion was extremely low. For detecting body condition changes, the flash seems to be harmful, probably by illuminating the lynx from the side, thus suppressing the lateral shadows that reveal bony contours, like the points of the ischium (hips) or the hollow of the flank.

Our study aimed at using population monitoring images for early detection of clinical signs, as a complement to other event-based surveillance systems. We observed that the diversity of conditions in which the images are captured allow the detection of a variety of clinical signs (Lenglin et al., submitted 2025). The database of monitoring images is also an interesting source for retrospective studies. However, our observations mean also that for quantifying the prevalence or characterizing different forms of the disease, an optimization of photographic conditions is necessary for each kind of change, e.g. nocturnal for skin changes and diurnal for body condition alterations. Therefore, to monitor specifically one kind of change, it is indispensable to consider technical improvements like maintaining a short shutter speed to avoid flash, to survey body condition changes. It could be useful to have different camera traps with different angles/flash conditions in a same place to monitor simultaneously different clinical changes.

Taking image quality into account can also help to refine predictive models for the incidence of a disease in wild populations, by considering doubtful images as potential positives with lower statistical weights, as in Murray et al. 2021).

The detectability of disease however can also be affected by behavior modifications. For example, Borchard, Eldridge, et Wright (2012) observed that wombats, that are habitually nocturnal, became diurnal when affected by sarcoptic mange. For changes where the detectability differs according to the time of day, this may constitute a bias of over-detection of severe cases. So even for the surveillance of one specific disease, a diversity of conditions of capture can be useful.

The study carried out here has enabled us to show that using camera traps to monitor the health status of an elusive species such as the lynx is relevant. In fact, it is quite possible to detect problems. However, the sensitivity (i.e., the ability to detect as many positive cases as possible) and the specificity (i.e., the ability to detect as few false positives as possible) are highly dependent on the protocol used. In this exploratory study, we found that the detection of different types of disorders was associated with different co-variables. A monitoring protocol such as that used to monitor the species by photo-identification therefore represents an opportunity to monitor different types of disorders without preconception. Conversely, in order to monitor a specific type of disorder, it is necessary to choose the appropriate shooting context. Our study is a retrospective exploratory study that will enable us to discern trends, and should therefore be supplemented by others that will enable to explore in greater detail the effect of the shooting context for a given type of disorder.

## Acknowledgments

The authors would like to thank all those involved in the Loup-Lynx network who collected and made available the data used in this study. We would also like to thank the external contributors who provided advice and expertise, in particular Philippe Gourlay and Etienne Levy.

