## Supplemental Tables 1, 2 and 3 for "Annotating opportunistic camera-trap images with conditions of recording, for the disease surveillance of Eurasian lynx (*Lynx lynx*)"

### *Supplementary Material*

Table 1. Estimation of the lynx visible part to calculate the distance estimator

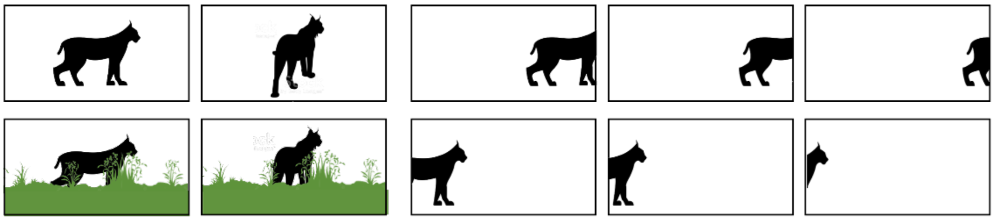

| | | | | |
|---|---|---|---|---|
| ***Lynx visible part*** | **1** | **0,75** | **0,5** | **0,25** |

Table 2. Estimation of the lynx angle: calculation of the *Profile* and *Profile_completeness* variables

| **Lynx visible part**<br>*Each part must be perfectly visible* | **Illustration**<br>*Visible part of the lynx is in orange* | **Profile** | **Profile_completeness** |
|---|---|---|---|
| *Right face* **AND** *Left face* | 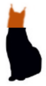 | **Face** | ***0.25*** |
| *Back of right ear***AND** *Back of left ear* | 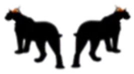 | **Back** | ***0.25*** |
| *Right face **AND** Right shoulder **AND** Right flank **AND** Right hip* | 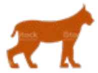 | **Right_Profile** | ***1*** |
| *Right face* **AND** *Right shoulder* **AND** *Right flank*<br>***OR***<br>*Right shoulder* **AND** *Right flank* **AND** *Right hip* | 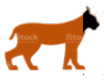 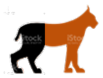 | | ***0.75*** |
| *Right face* **AND** *Right shoulder*<br>**OR**<br>*Right flank **AND** Right hip* | 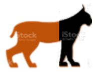 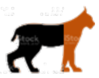 | | ***0.5*** |
| *Right face*<br>**OR**<br>*Right hip* | 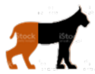 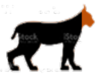 | | ***0.25*** |
| *Left face **AND** Left shoulder **AND** Left flank **AND** Left hip* | 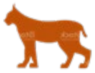 | **Left_Profile** | ***1*** |
| *Left face* **AND** *Left shoulder* **AND** *Left flank*<br>***OR***<br>*Left shoulder* **AND** *Left flank* **AND** *Left hip* | 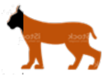 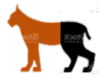 | | ***0.75*** |
| *Left face* **AND** *Left shoulder*<br>**OR**<br>*Left flank **AND** Left hip* | 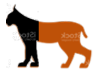 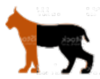 | | ***0.5*** |
| *Left face*<br>**OR**<br>*Left hip* | 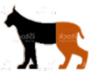 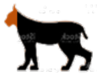 | | ***0.25*** |

Table 3. Pair-wise comparison estimators

| | | Cutaneous change detection | | | Body condition change detection | | |
|---|---|---|---|---|---|---|---|
| | | **Test performed** | **p-value** | **Significance (S) p-value < 0.05** | **Test performed** | **p-value** | **Significance (S) p-value < 0.05** |
| **Settings** | **Flash** | Chi-square | 0.76 | NS | Fisher | 0.03 | S |
| | **Colour range of the image** | Fisher | 0.30 | NS | Fisher | 0.05 | S |
| | **Exposure value** | Wilcoxon-Mann-Whitney | 0.08 | NS | Wilcoxon-Mann-Whitney | 0.25 | NS |
| | **Aperture of the diaphragm** | Wilcoxon-Mann-Whitney | 0.82 | NS | Wilcoxon-Mann-Whitney | 0.55 | NS |
| | **Diaphragm shutter speed** | Wilcoxon-Mann-Whitney | 0.08 | NS | Wilcoxon-Mann-Whitney | 0.54 | NS |
| | **ISO** | Wilcoxon-Mann-Whitney | 0.38 | NS | Wilcoxon-Mann-Whitney | 0.26 | NS |
| **Light conditions** | **Daytime/Nighttime** | Chi-square | 0.84 | NS | Fisher | 0.03 | S |
| **Lynx position** | **Right face** | Chi-square | 0.76 | NS | Chi-square | 0.04 | S |
| | **Right shoulder** | Chi-square | 0.82 | NS | Chi-square | 0.02 | S |
| | **Right flank** | Chi-square | 0.67 | NS | Fisher | 0.003 | S |
| | **Right hip** | Chi-square | 0.84 | NS | Chi-square | 0.002 | S |
| | **Back of right ear** | Chi-square | 0.95 | NS | Fisher | 0.06 | S |
| | **Tail** | Chi-square | 0.62 | NS | Chi-square | 0.81 | NS |
| | **Left face** | Chi-square | 0.07 | NS | Fisher | 0.003 | S |
| | **Left shoulder** | Chi-square | 0.05 | S | Fisher | 0.02 | S |
| | **Left flank** | Chi-square | 0.17 | NS | Fisher | 0.06 | NS |
| | **Left hip** | Chi-square | 0.19 | NS | Fisher | 0.01 | S |
| | **Back of left ear** | Chi-square | 0.71 | NS | Fisher | 0.69 | NS |
| | Length of cropped image on lynx/length of entire image | Wilcoxon-Mann-Whitney | 0.95 | NS | Wilcoxon-Mann-Whitney | 0.43 | NS |
| | Height of cropped image on lynx/height of entire image | Wilcoxon-Mann-Whitney | 0.51 | NS | Wilcoxon-Mann-Whitney | 0.46 | NS |
| **Artifacts** | **Blur** | Chi-square | 0.02 | S | Fisher | 0.28 | NS |
| | **Light artifact** | Chi-square | 1 | NS | Fisher | 1 | NS |
| | **Environmental artifact** | Chi-square | 0.62 | NS | Fisher | 0.39 | NS |
| | **Weather artifact** | Fisher | 1 | NS | Fisher | 1 | NS |
| | **Definition of image length in pixels** | Wilcoxon-Mann-Whitney | 0.19 | NS | Wilcoxon-Mann-Whitney | 0.18 | NS |
| | **Definition of image heigth in pixels** | Wilcoxon-Mann-Whitney | 0.34 | NS | Wilcoxon-Mann-Whitney | 0.25 | NS |
